# Lipid loss and compositional change during preparation of liposomes by common biophysical methods

**DOI:** 10.1101/2024.05.30.596670

**Authors:** Eunice Kim, Olivia Graceffa, Rachel Broweleit, Ali Ladha, Andrew Boies, Robert J. Rawle

**Author notes:** **<u>Corresponding Author Contact Information</u>**.

## Abstract

Liposomes are widely used as model lipid membrane platforms in many fields, ranging from basic biophysical studies to drug delivery and biotechnology applications. Various methods exist to prepare liposomes, but common procedures include thin-film hydration followed by extrusion, freeze-thaw, and/or sonication. These procedures have the potential to produce liposomes at specific concentrations and membrane compositions, and researchers often assume that the concentration and composition of their liposomes are similar to, if not identical, to what would be expected if no lipid loss occurred during preparation. However, lipid loss and concomitant biasing of lipid composition can in principle occur at any preparation step due to nonideal mixing, lipid-surface interactions, etc. Here, we report a straightforward method using HPLC-ELSD to quantify the lipid concentration and membrane composition of liposomes, and apply that method to study the preparation of simple POPC/cholesterol liposomes. We examine many common steps in liposome formation, including vortexing during re-suspension, hydration of the lipid film, extrusion, freeze-thaw, sonication, and the percentage of cholesterol in the starting mixture. We found that the resuspension step can play an outsized role in determining the overall lipid loss (up to ∼50% under seemingly rigorous procedures). The extrusion step yielded smaller lipid losses (∼10-20%). Freeze-thaw and sonication could both be employed to improve lipid yields. Hydration times up to 60 minutes and increasing cholesterol concentrations up to 50 mole% had little influence on lipid recovery. Fortunately, even conditions with large lipid loss did not substantially influence the target membrane composition more than ∼5% under the conditions we tested. From our results, we identify best practices for producing maximum levels of lipid recovery and minimal changes to lipid composition during liposome preparation protocols. We expect our results can be leveraged for improved preparation of model membranes by researchers in many fields.

**Statement of Significance:** Liposomes are spherical lipid membranes that can be prepared by a variety of biophysical techniques. Researchers use liposomes in a variety of ways, including fundamental biophysical studies of lipid membranes, in drug delivery, drug formulation, and other biotechnology applications. In this report, we study the process to prepare liposomes by several common techniques and validate how reliable each technique is at producing consistent liposome concentrations and lipid compositions. We identify best practices for researchers to produce reliable liposome preparations.

## 1. Introduction

Liposomes are model lipid membranes widely used as cell membrane mimics (1–8) in both applied and basic scientific studies by many laboratories, including ours (9, 10). They can be prepared in a variety of size ranges, including small unilamellar vesicles (SUVs, diameter (d) < ∼100 nm), large unilamellar vesicles (LUVs, ∼100 nm < d < ∼1 µm) or giant unilamellar vesicles (GUVs, diameter > ∼1 µm). In this report, we will focus exclusively on SUVs and LUVs.

Various procedures exist to prepare SUVs and LUVs (11–18). A common process in many protocols is to combine purified lipid components in the desired ratio in organic solvent, dry to a lipid film under an inert gas, and then re-suspend in water or buffer to form a multilamellar lipid suspension. This suspension then undergoes one of several liposome formation techniques, the most common of which are extrusion, freeze-thaw, or sonication, although others exist as well (14–17). Extrusion involves passing the multilamellar lipid suspension through a track-etched polycarbonate membrane of defined pore size (typically 30-200 nm) to produce SUVs and/or LUVs of a diameter similar to the pore size. Freeze-thaw procedures consist of several freeze-thaw cycles of the re-suspended lipid film, often followed by extrusion. These freeze-thaw cycles are typically included to improve liposome encapsulation efficiency of molecular cargo loaded into the re-suspension buffer (19). Sonication procedures use either bath or probe tip sonication to break down the re-suspended multilamellar structures into liposomes. Sonication is procedurally faster and more straightforward than extrusion, but with less reproducibility in its liposome size distributions (20, 21).

In principle, each of these liposome preparation procedures allow the experimenter to prepare SUVs and/or LUVs at a desired concentration and membrane lipid composition, hence their widespread usage in many fields. In practice however, it is difficult and/or tedious to verify the lipid concentration and composition of the resulting liposomes, and so researchers often assume that the lipid concentration and composition of their liposomes are similar to, if not identical, to what would be expected if no lipid loss occurred during any of the preparation steps. Indeed, standardized measurements of lipid loss and compositional variance are not commonly reported for liposome preparations in the literature.

However, limited evidence suggests that lipid composition may vary both within a sample (22–24) and between samples prepared differently (25, 26). For example, cholesterol solubility in liposome membranes has been reported to vary substantially depending on preparation method (26). In principle, lipid loss and biasing of lipid composition could occur at any step during preparation due to nonideal mixing, lipid-surface interactions, etc. Therefore, in this report we study the extent of lipid recovery and average compositional variability in liposomes prepared by extrusion, freeze-thaw, or sonication.

Inspired by methods to study liposome-based nutritional supplements and drug formulations (27–31), we have developed a method to quantify the lipid composition of liposomes using high-performance liquid chromatography with evaporative light scattering detection (HPLC-ELSD). Peak identification in our HPLC-ELSD approach is relatively straightforward since the lipid components are known a priori, and the retention time of each peak can be compared with a standard. Similarly, quantification of each lipid component is achieved with a standard curve. Crucially, the standard curve for each sample set is prepared from the same starting lipid mixture as the samples to be measured, such that any changes in lipid recovery and composition can be unambiguously identified. We note however that this approach only allows us to assess lipid loss and average compositional changes for liposomes at the bulk level; variation between individual liposomes is a separate and important metric, but is not possible to be addressed by our methods.

Using our HPLC-ELSD method, we studied the preparation of liposomes composed of simple POPC/cholesterol lipid mixtures, a commonly used 2-component model membrane lipid composition. We examined the influence of many common steps in liposome formation procedures on both the resulting lipid recovery and compositional variability, including vortexing during re-suspension, hydration “swelling” of the lipid film, extrusion, freeze-thaw, bath sonication, and the percentage of cholesterol in the starting mixture during liposome preparation. Our goal with this report is to identify best practices for producing maximum levels of lipid recovery and minimal changes to the lipid composition during liposome formation protocols. We therefore expect our results can be leveraged for improved preparation of model membranes in many fields.

## 2. Materials and Methods

### 2.1. Materials

Palmitoyl oleoyl phosphatidylcholine (POPC) and cholesterol were obtained from Avanti Polar Lipids (Alabaster, AL). Chemical reagents such as ethanol, methanol, chloroform, and ammonium formate were obtained from Thermo Fisher Scientific (Waltham, MA) and Sigma Aldrich (St. Louis, MO). Milli-Q H2O was purified using a ThermoScientific Millipore Milli-Q lab water system. Vortexers used included the Vortex Genie (Scientific Industries Inc, Bohemia, NY), the Fisherbrand™ Analog Vortex Mixer (Thermo Fisher Scientific, Waltham, MA, cat # 02-215-414), and the BV1000 Vortex Mixer (Benchmark Scientific, Edison, NJ).

### 2.2. Preparation of Lipid Mixtures and the Post-Resuspension Solution

Isolated lipid stock solutions consisted of 10g/L cholesterol and 25g/L POPC, both dissolved in chloroform, and were stored at -20°C. Lipid stock solutions in organic solvent were handled using Hamilton syringes (1700 series, Hamilton Company, Franklin, MA) that were solvent compatible with both chloroform and methanol.

Lipid mixtures were prepared by adding the appropriate volumes of lipid stock solutions to a glass autosampler vial, which was kept capped as much as possible during preparation to minimize evaporation. This is referred to as the Parent Lipid Mixture. For reference, this process, as well as those described below, is represented in a schematic in most figures, such as Figure 1A.

**Figure 1.**
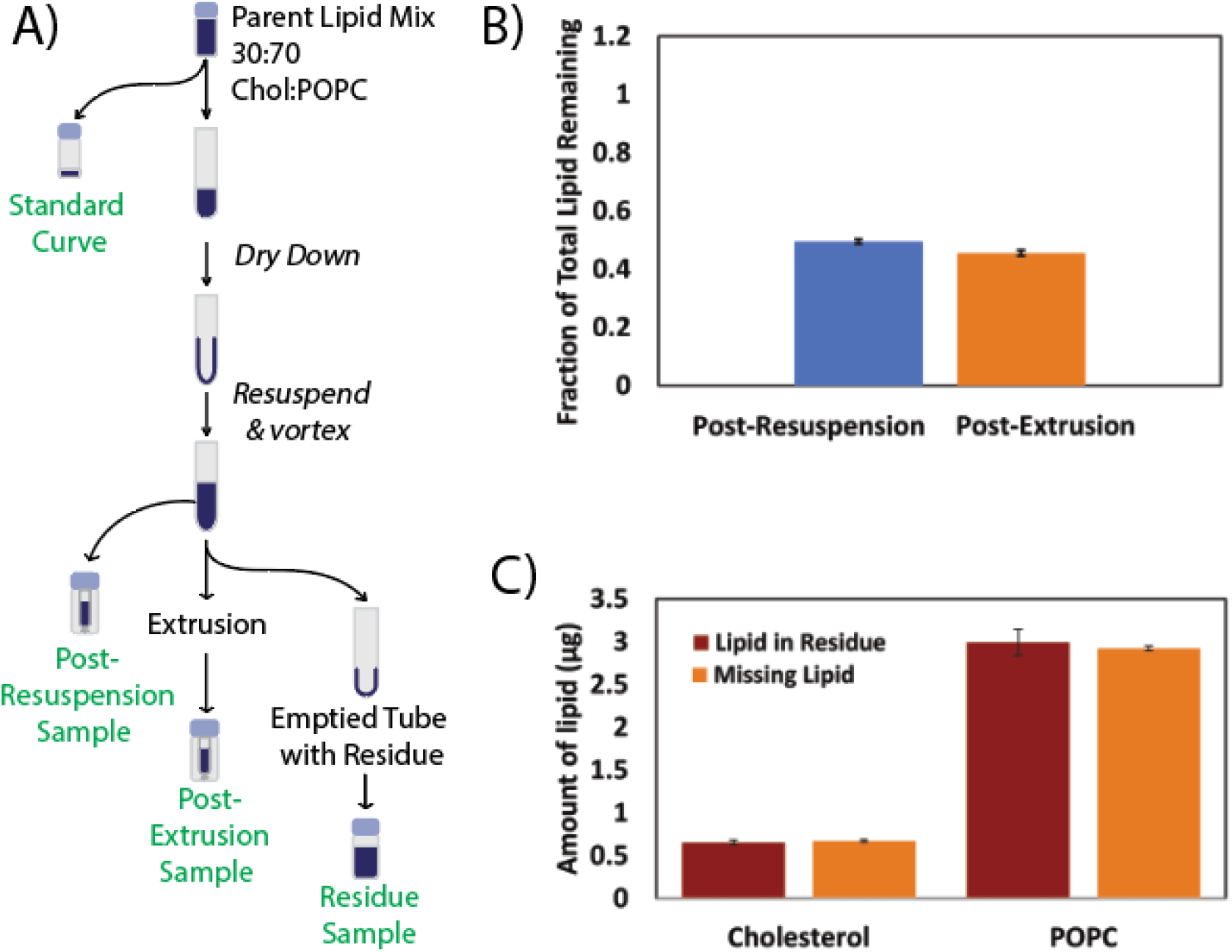
Standard extrusion procedure can yield substantial lipid loss. **A)** shows a schematic of liposome preparation and sample collection for HPLC-ELSD analysis. Liposomes were prepared by the Standard Extrusion Protocol from 30:70 cholesterol: POPC lipid mixtures following the procedures listed in the Materials and Methods. The standard curve was prepared directly from the parent lipid mixture. The post-resuspension sample was taken immediately following the resuspension of the dried lipid film, but prior to extrusion. The Vortex-Genie (30 seconds at max speed) was used for resuspension. The post-extrusion sample was taken immediately following the extrusion step. Following collection, all samples (green text) were analyzed by HPLC-ELSD. **B)** shows the fraction of total lipid remaining for each sample, calculated relative to the theoretical maximum moles of lipid that could be recovered if the entirety of the starting lipid material ended up in that sample (see Equation 1). Data shown is mean ± propagated error of 3 measurements (see Equation 2 for error calculations). **C)** shows the amount of lipid recovered in the “residue” remaining in the emptied test, compared to the calculated amount of missing lipid. Missing lipid was calculated by subtracting the amount recovered in the post-resuspension sample from the starting lipid amount. Data shown is mean ± stdev of 3 measurements

A portion of this Parent Lipid Mixture was removed to a separate vial, ultimately used to prepare the standard curve. This is referred to as the Standard Curve sample. The remainder of the Parent Lipid Mixture was then separated into glass test tubes for liposome synthesis. All samples were then dried while rotating under nitrogen gas to evaporate the organic solvent, followed by vacuum desiccation for 2-24 hours. This left a lipid film at the bottom of each vessel.

To hydrate and re-suspend the dried lipid films intended for liposome synthesis, Milli-Q H_2_O (typically 270 µL) was added to each test tube, incubated at room temperature for a hydration period (typically 15 minutes, although this time was tested in some experiments as listed), and then vortexed vigorously (typically 30 seconds at maximum speed, although this was varied in some experiments as listed). This cloudy re-suspended solution is referred to as the Post-Resuspension solution.

### 2.3. Preparation of the Standard Curve sample for HPLC-ELSD analysis

Following removal from the Parent Lipid Mixture and after being dried to a lipid film as described above, the Standard Curve sample was then dissolved in HPLC-grade methanol to the desired concentration. This solution was capped and sealed with Teflon tape to prevent evaporation and stored at -20°C until analysis by HPLC-ELSD.

### 2.4. Liposome Preparation Method 1: Standard Extrusion Protocol

To prepare liposomes by the Standard Extrusion protocol, the Post-Resuspension solution was extruded through a 100 nm pore size polycarbonate membrane for twenty passes using an Avanti mini-extruder (Avanti Polar Lipids, Alabaster, AL), following the manufacturer’s instructions.

### 2.5. Liposome Preparation Method 2: Freeze-Thaw Extrusion

Liposomes prepared by freeze-thaw extrusion underwent a similar procedure as the Standard Extrusion protocol, but with an additional step at the beginning. First, the Post-Resuspension solution in its glass test tube was rapidly frozen with liquid nitrogen (−196°C) for 5 minutes and then thawed with a lukewarm water bath (30°C) for 5 minutes. This freeze-thaw cycle was conducted 5 times. After the freeze-thaw cycles, the Standard Extrusion protocol was followed.

### 2.6. Liposome Preparation Method 3: Sonication

To prepare liposomes by sonication, the Post-Resuspension solution in its glass test tube was sonicated in a bath sonicator (CPXH series, Branson Ultrasonics, CT) for 1 hour at room temperature. After the 1 hour sonication, the cloudy suspension was observed to become clear, indicative of liposome formation.

### 2.7. Sampling the “Residue” for HPLC-ELSD analysis

Residue samples were collected to quantify the amount of residual lipid coating the glass test tube following re-suspension of the dried lipid film. After emptying the test tube of the Post-Resuspension solution, a Kimwipe was folded in thirds lengthwise and twisted to plug the test tube. The test tube was placed in a benchtop centrifuge (Fisher Scientific, Model 228) upside down and spun at the highest speed for a few seconds. This was done to remove any remaining drops of solution in the test tube. 500µL of HPLC-grade methanol was then added down the sides of the test tube. The test tube was vortexed for 30 seconds and then transferred to an autosampler vial. The vial was immediately capped and sealed with Teflon tape to prevent evaporation and stored at -20°C until analysis by HPLC-ELSD.

### 2.8. Sampling the Post-Resuspension and Liposome solutions for HPLC-ELSD analysis

To prepare HPLC samples from any stage of liposome preparation following re-suspension of the dried lipid film in H_2_O (e.g. the Post-Resuspension solution, after liposome formation, etc.), a small volume of the aqueous suspension (typically 12uL) was removed by pipette from the top of the liquid into an autosampler vial, and extensively dried under a steady stream of nitrogen gas, followed by vacuum desiccation for 2-4 hours to remove any residual H_2_O. The resulting lipid film was then completely dissolved in methanol (typically 240uL), vortexed for 30 seconds, and then transferred to a glass vial insert with polyspring. This vial was capped and sealed with Teflon tape to prevent evaporation and stored at -20°C until analysis by HPLC-ELSD.

### 2.9. HPLC-ELSD Method

High performance liquid chromatography (HPLC) (Agilent Technologies 1200 Series) with an autosampler injector, evaporative light scattering detection (ELSD) (Agilent Technologies 1260 Series), operated using ChemStation software, was used to separate and quantify lipids. An isocratic method was employed, with a mobile phase of 10mM ammonium formate in methanol, filtered by vacuum glass filtration with Durapore 0.22µm PVDF membrane filter. This method has been previously established (32) to work well for the lipids tested herein. HPLC settings: liquid flow rate = 0.500mL/min, column temperature = 35°C. ELSD settings: temperature = 70°C, gain = 10, nitrogen gas pressure = 3.5 bar.

Standard curves were collected by injecting different volumes (typically 9-27 µL) of the standard curve sample, and integrating the resulting ELSD peaks. Each standard curve injection was collected in triplicate, and was collected within 1-2 days of the experimental samples. The injection volume of experimental samples varied from 5-50 µL depending on the lipid prep, to ensure that each measurement fell within the standard curve.

### 2.10. HPLC-ELSD Data Analysis and Error Calculations

Data analysis of individual peaks in the chromatograms was performed using ChemStation software. The ELSD response with respect to concentration of the analyte is nonlinear, as has been established (33). Therefore, integrated peak areas for experimental samples were converted into amount of lipid detected (either mass or moles) by using interpolation of adjacent measurements on the standard curve.

In data figures throughout this report, fraction of lipid remaining (*Fract*_*TotLipRem*_) was calculated as:

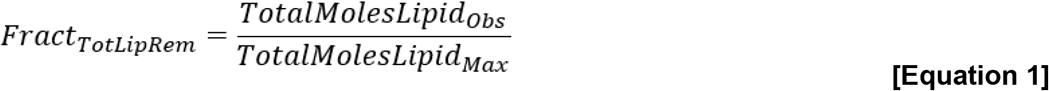

where *TotalMolesLipid*_*Obs*_ refers to the sum of moles of all lipids measured to be in the sample by HPLC-ELSD analysis using the standard curve. *TotalMolesLipid*_*Max*_ was defined as the maximum moles of lipid that could theoretically be recovered in a given sample if the entirety of the starting material ended up in that sample. This theoretical maximum number was calculated from the molar concentration of lipids in the Parent Lipid Mixture and the volume of the Parent Lipid Mixture used to prepare each sample.

Propagated error for the fraction of total lipid remaining relative to the theoretical maximum (*Error*_*FractLipRem*_) was calculated as:

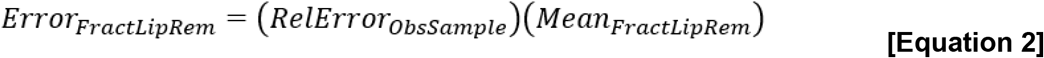

where *Mean*_*FractLipRem*_ was the mean fraction of total lipid remaining, and *RelError*_*ObsSample*_ was the propagated relative error of the observed total lipid in the sample, calculated as

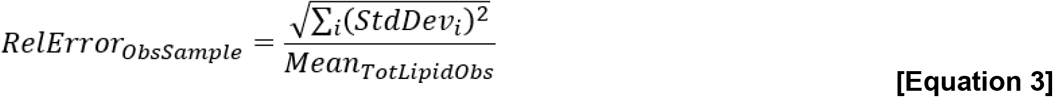

where *Mean*_*TotLipidObs*_ is the mean moles of total lipid observed in the sample, and *StdDev*_*i*_ is the standard deviation of the observed moles of the *ith* lipid, determined from at least 3 replicate measurements.

Mole fraction of each lipid was calculated according to the standard definition: observed moles of the lipid of interest divided by the total observed moles of all lipids.

### 2.11. Dynamic Light Scattering (DLS)

DLS was used to determine the size distribution of liposomes prepared by the different liposome preparation methods. Following preparation, each liposome suspension was diluted to 0.1 µg/mL total lipid concentration using MilliQ H_2_O. All MilliQ H_2_O used for dilution was centrifuged briefly using a tabletop mini-centrifuge to remove any large dust particles. 100 µL of each sample was analyzed using a DynaPro NanoStar DLS (Wyatt Technologies, Santa Barbara, CA) with temperature control set at 20°C. 10 acquisitions were averaged, and analyzed using the Hollow Sphere model in the DYNAMICS software package (Wyatt Technologies).

## 3. Results and Discussion

### 3.1. Overview of lipid quantification by HPLC-ELSD

We used HPLC-ELSD to quantify lipid recovery and compositional changes at different steps of each liposome preparation procedure (see Materials and Methods). It is important to note that each experiment was designed such that a comparison sample was extracted from the lipid mixture in organic solvent prior to any other manipulation (see schematic associated with each data figure, such as **Figure 1a**). This comparison sample was used to prepare the standard curve for that data set. This approach allowed for unambiguous quantification of lipid recovery and compositional changes during liposome preparation.

### 3.2. Standard extrusion protocol can produce substantial lipid loss

As a test case, we utilized our HPLC-ELSD approach to examine lipid loss and compositional bias that might occur during our “standard extrusion protocol” (**Figure 1a**). This standard extrusion protocol (see Materials and Methods) is characteristic of extrusion procedures employed by many labs, and includes preparation of the lipid mixture in organic solvent, thorough evaporation of the solvent which leaves a lipid film, re-suspension of the lipid film in water by vigorous vortexing for 30 seconds, followed by extrusion. In our initial data, we prepared liposomes composed of a binary mixture of 30:70 mol% cholesterol:POPC, two lipids commonly employed in model membrane studies. This served as our test case lipid composition for much of the data in this report.

In initial data sets, we observed that lipid loss could be quite substantial. When we compared the total lipid content of the liposomes post-extrusion to the parent lipid mixture, we found that up to 50-60% of the total lipid had been lost (**Figure 1b**). The bulk of this lipid loss occurred during the re-suspension of the lipid film prior to extrusion, despite vigorous vortexing during resuspension. However, this was confirmed by using methanol to extract the lipid residue which remained in the glass test tube following resuspension, and quantifying by HPLC-ELSD (**Figure 1c**). We observed that the entirety of the “missing” lipid could be recovered from this lipid residue. The extrusion step itself resulted in the loss of only a small portion (∼5%) of the total lipid (or ∼10% of the lipid remaining after the resuspension step).

If not identified or corrected, such a substantial loss of lipid during liposome preparation could easily lead to problematic experimental results, especially for applications which are sensitive to liposome or lipid concentration. Additionally, if one or more of the lipid components is selectively depleted relative to the other during this lipid loss, the average lipid composition itself could be substantially altered. Below, we investigate each of these issues, and identify best practices to minimize lipid loss.

### 3.3. Importance of vortex step on lipid loss and compositional variance

Given that re-suspension of the lipid film from the glass test tube was the step that produced the greatest loss of lipid, we examined the role that vortexing could play in that step. We tested the brand/model of vortex machine, the vortex method, the time of vortexing, and the speed of vortexing. In each comparison, we used our test case composition of 70/30 mol% POPC/cholesterol, and re-suspended via vortexing following a hydration step of 15 minutes at RT. We then quantified lipid recovery after re-suspension and the resulting average lipid composition using our HPLC-ELSD method.

We observed that the brand/model of vortex machine had an outsized role in determining the amount of lipid loss (**Figure 2**). We compared 3 different brands/models (Vortex-Genie from Scientific Industries, BV1000 from Benchmark Scientific, and the Fisherbrand™ Analog Vortex Mixer), vigorously vortexing the sample for 30 seconds at the maximum speed (3200 rpm for each model). These different vortexers represent older vs newer models that might be available in different research laboratories. We observed considerable variability between brands/models. Both the Fisherbrand™ and BV1000 vortexers produced nearly complete recovery during re-suspension under these vortex conditions, while the older Vortex-Genie brand consistently yielded poor recovery with a rather wide range (∼40-70% at 30 seconds at maximum speed). Fortunately however, the poor recovery of the Vortex-Genie brand during re-suspension did not result in substantial alteration of the resulting average lipid composition from the target composition, at least for the simple test case composition of 70/30 mol% POPC/cholesterol (**Figure 2**).

**Figure 2.**
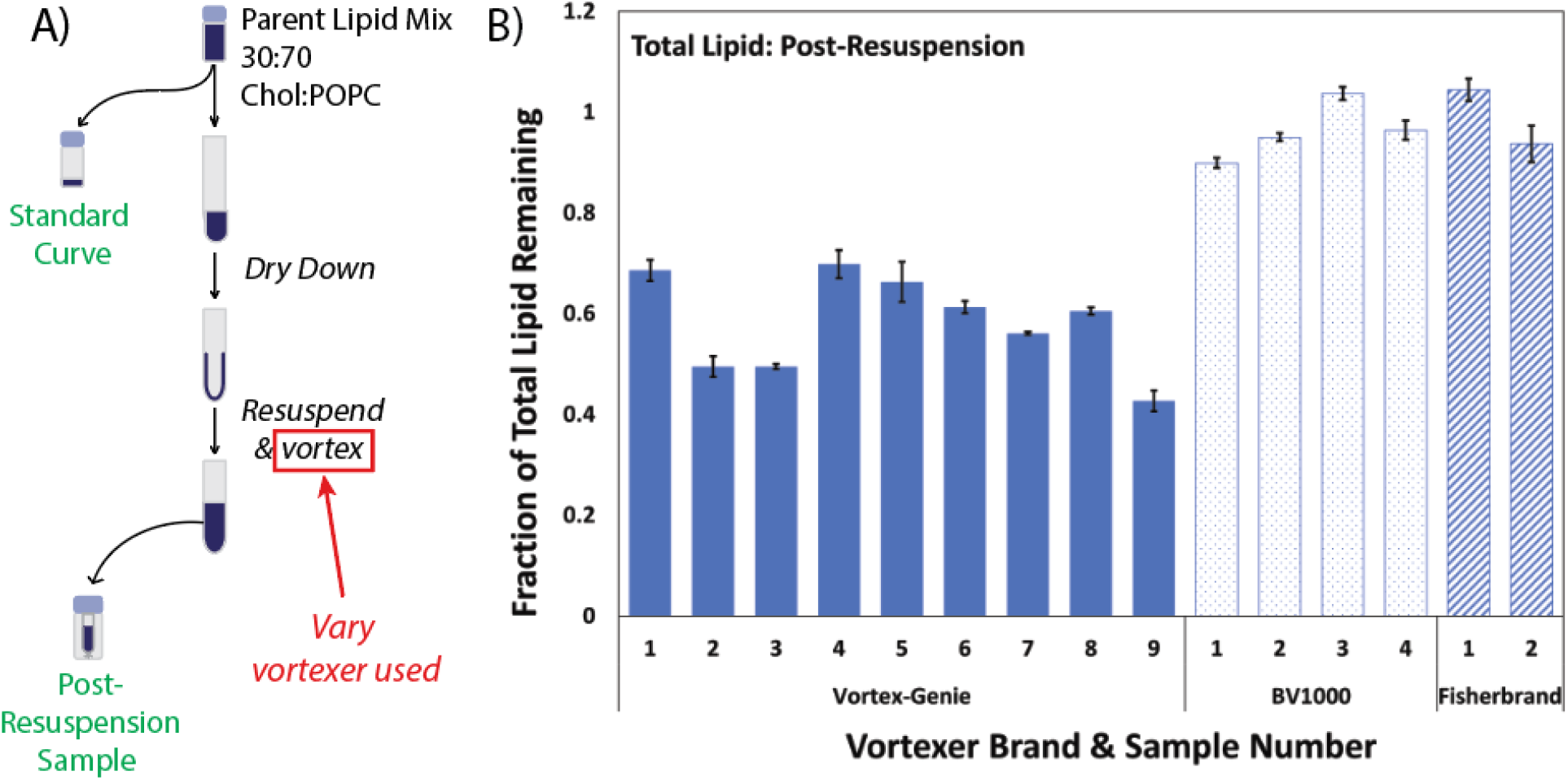
Fraction of total lipid remaining for post-resuspension samples prepared with different vortexers (Vortex-Genie, BV1000, Fisherbrand™). **A)** shows a schematic of sample preparation. Post-resuspension samples were prepared from 30:70 cholesterol: POPC lipid mixtures following the procedure listed in the Materials and Methods, using the vortexer brand as listed (30 sec at max speed). Following sample collection, all samples were analyzed by HPLC-ELSD. **B)** shows the fraction of total lipid remaining for each sample, calculated relative to the theoretical maximum moles of lipid that could be recovered if the entirety of the starting lipid material ended up in that sample (see Equation 1). Data shown is mean ± propagated error of 3 measurements (see Equation 2 for error calculations).

**Figure 3.**
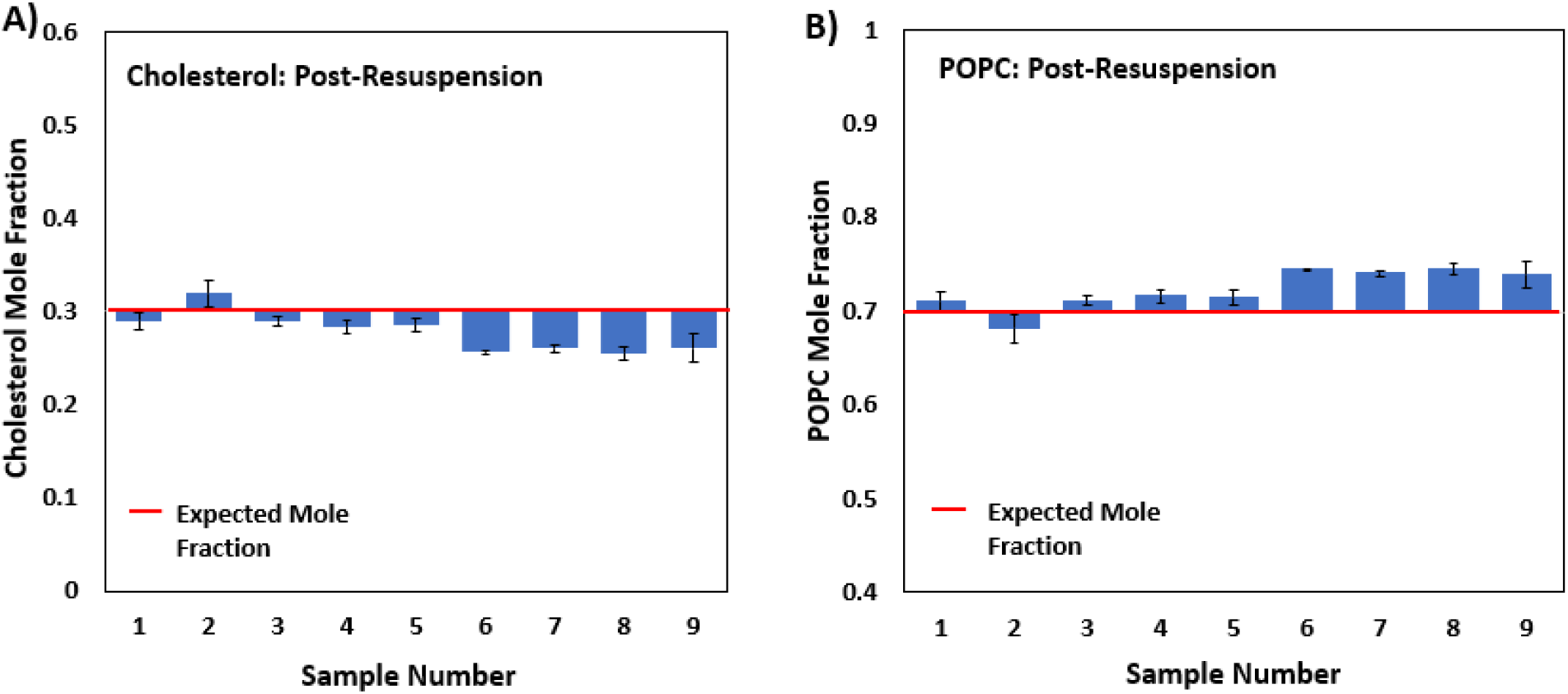
Measured mole fractions of A) cholesterol and B) POPC for post-resuspension samples prepared using the Vortex-Genie. Sample numbers correspond to those in Figure 2B above. The expected mole fraction (red line) represents the mole fraction in the Parent Lipid Mixture prior to any manipulation. For each sample, data shown is the mean ± stdev of 3 replicate measurements.

In our experience, a 30 sec vortex window is often longer than many researchers typically employ when vortexing samples. Therefore, we also tested the effect of vortex time on the extent of lipid loss, using the Fisherbrand™ vortexer. We tested vortex times of 0, 10, and 30 sec following hydration (**Figure S1A**). As expected, lipid recovery increased with vortexing time. We observed that while maximum lipid recovery was reliably achieved by 30 seconds, at 10 seconds most of the lipid had been recovered. As before, even when we observed lower lipid recovery at 10 seconds, we found that the resulting average lipid composition was not substantially altered from the target composition of 70/30 mol% POPC/cholesterol.

We also examined the effect of vortex speed on lipid loss using the Fisherbrand™ vortexer, testing Speeds 4, 7, and 10 out of 10 (corresponding to 3200 rpm). To best illustrate the effect of vortex speed, we employed a vortex time of 10 seconds, which yielded nearly complete lipid recovery at the maximum vortex speed (**Figure S1B**), but which would presumably demonstrate sensitivity to lower vortex speeds. Perhaps predictably, we observed that as vortex speed decreased, we observed less lipid recovery, again underscoring the need for researchers to employ lengthy vortex procedures at maximum speed to ensure reliable and maximum recovery. Fortunately however, the resulting average lipid composition remained close to the target composition for all samples with at least some lipid recovery.

We also note that for nearly all of the poor recovery samples, there was little to no evidence of a lipid residue in the test tube that was visible by eye, indicating that simply re-suspending until the lipid film appeared to be gone by visual inspection would not be a reliable strategy.

### 3.4. Influence of hydration time on lipid loss and composition

Thin-film hydration is a procedure in which the re-suspension liquid is added to the dried lipid film and incubated before vortexing or other resuspension of the film (34). This hydration technique can swell the lipids on the surface of the test tube (18). If the lipid film is swelled in such a manner, it may detach more easily from the test tube upon vortexing, thereby increasing the lipid recovery during re-suspension. To test whether thin-film hydration increases lipid recovery following vortexing, we hydrated our dried lipid films in MilliQ water for 0 minutes, 15 minutes and 60 minutes at room temperature, vortexed for 30 seconds at maximum speed using the Vortex-Genie (our worst model), and then quantified lipid recovery and composition by HPLC-ELSD (**Figure 4**). We found that there was little difference in lipid recovery between the different hydration periods we tested (**Figure 4b**), suggesting that thin-film hydration does not have a large influence on lipid recovery during liposome preparation under these conditions. Once again, the resulting lipid composition for each hydration condition showed only minor variations from baseline (**Figure 4c and d**). Note that these results do not preclude the possibility that hydration may influence other important variables, such as the extent of unilamellarity, or encapsulation efficiency, which we did not study here.

**Figure 4.**
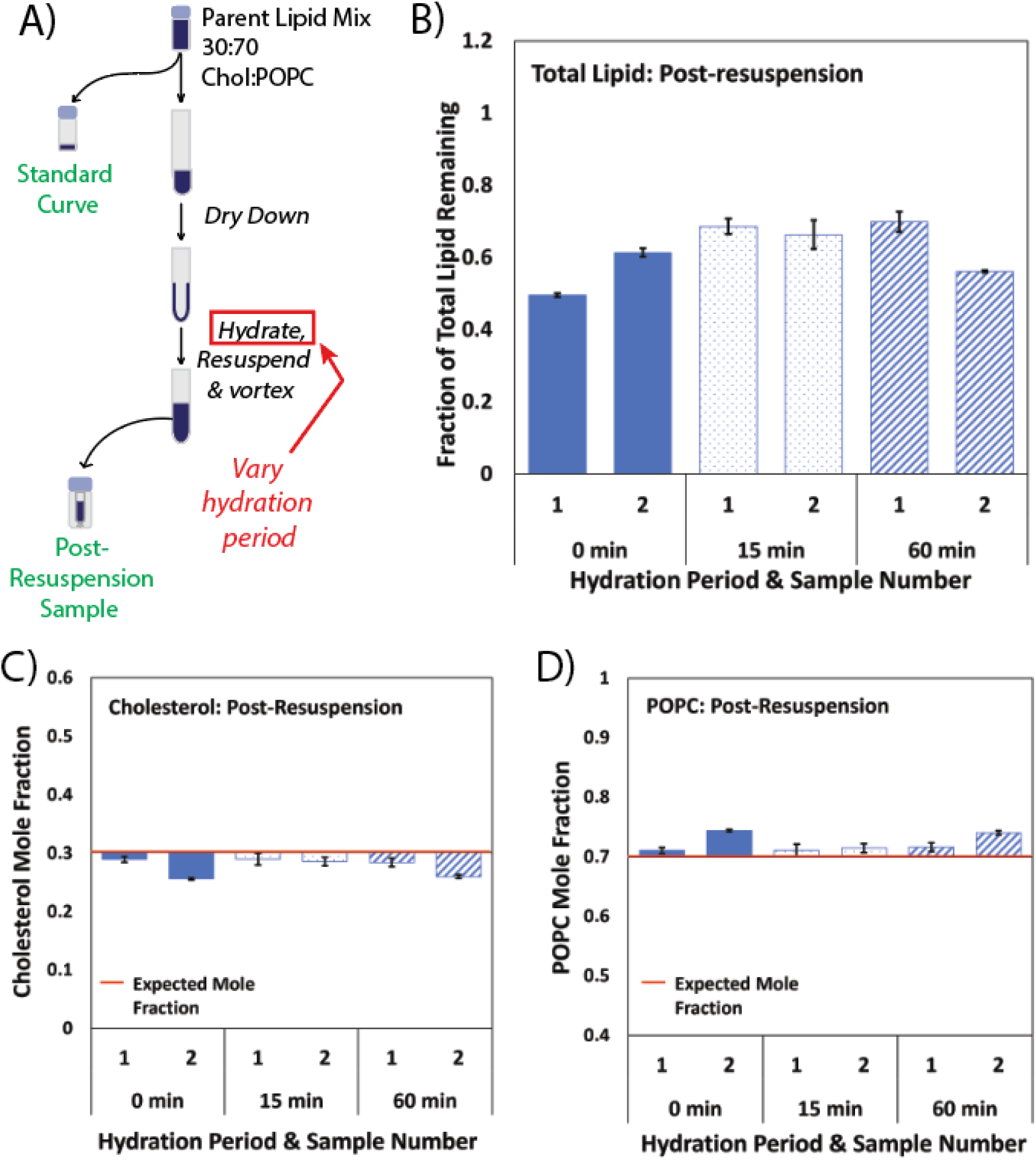
Different hydration periods prior to re-suspension yield little difference in lipid recovery or resulting lipid compositions. **A)** shows a schematic of sample preparation and collection for HPLC-ELSD analysis. Post-resuspension samples were prepared from 30:70 cholesterol: POPC lipid mixtures following the procedures listed in the Materials and Methods, but with variable hydration times (0 min, 15 min, or 60 min) prior to vortexing. The Vortex Genie (30 seconds at max speed) was used to resuspend all samples. Following sample collection, all samples were analyzed by HPLC-ELSD. **B)** shows the fraction of total lipid remaining for each sample, calculated relative to the theoretical maximum moles of lipid that could be recovered if the entirety of the starting lipid material ended up in that sample (see Equation 1). Data shown is mean ± propagated error of 3 measurements (see Equation 2 for error calculations). **C and D)** show the measured mole fractions of cholesterol and POPC for each sample. The expected mole fraction (red line) represents the mole fraction in the Parent Lipid Mixture prior to any manipulation. For each sample, data shown is the mean ± stdev of 3 replicate measurements.

### 3.5. Extrusion step itself typically produces minimal lipid loss

In our initial data (**Figure 1** above), the extrusion step was not the dominant source of lipid loss, however a small portion of the total lipid was lost during that step. To more fully quantify the extent and variability of lipid loss due to this step, we calculated the percent loss solely due to the extrusion step by comparing the quantified total lipid immediately before and immediately after extrusion for many samples of our test case composition (70% POPC, 30% cholesterol). We observed some variability, typically ranging from 0-20% total lipid loss, with an average of 7% (N = 12). Given the relatively small loss of lipid, it was not surprising that in our data the impact on lipid composition due to this step was minimal.

### 3.6. Sonication and freeze-thaw methods

To examine the influence of other liposome synthesis methods on lipid recovery and lipid composition, we performed a side-by-side comparison of 2 common methods of liposome synthesis in addition to the Standard Extrusion procedure – Freeze-Thaw Extrusion and Bath Sonication. We used our HPLC-ELSD method to quantify differences in lipid recovery and composition of the resulting liposomes produced by these 3 different methods (see **Figure 5**). Importantly, all samples were prepared from the same initial lipid mixture (see schematic in **Figure 5a**), such that any observed variation in the lipid recovery and/or lipid composition would only be due to differences in the liposome synthesis methods themselves. For the re-suspension step in each procedure, we used the Vortex Genie brand vortexer (consistently our worst, see **Figure 2** above) in order to better observe the effect of the liposome synthesis methods. Not surprisingly, we observed that the Bath Sonication procedure yielded essentially complete lipid recovery in the liposomes, indicating that sonication successfully removed any residual lipid film on the glass test tube following re-suspension by vortexing. The Freeze-Thaw Extrusion procedure yielded a higher lipid recovery (∼70%) than the Standard Extrusion procedure (∼50%), indicating that the multiple freeze-thaw cycles also improved recovery of the lipid film. All 3 methods did not produce any substantial change in the lipid composition from the starting lipid mixture.

**Figure 5.**
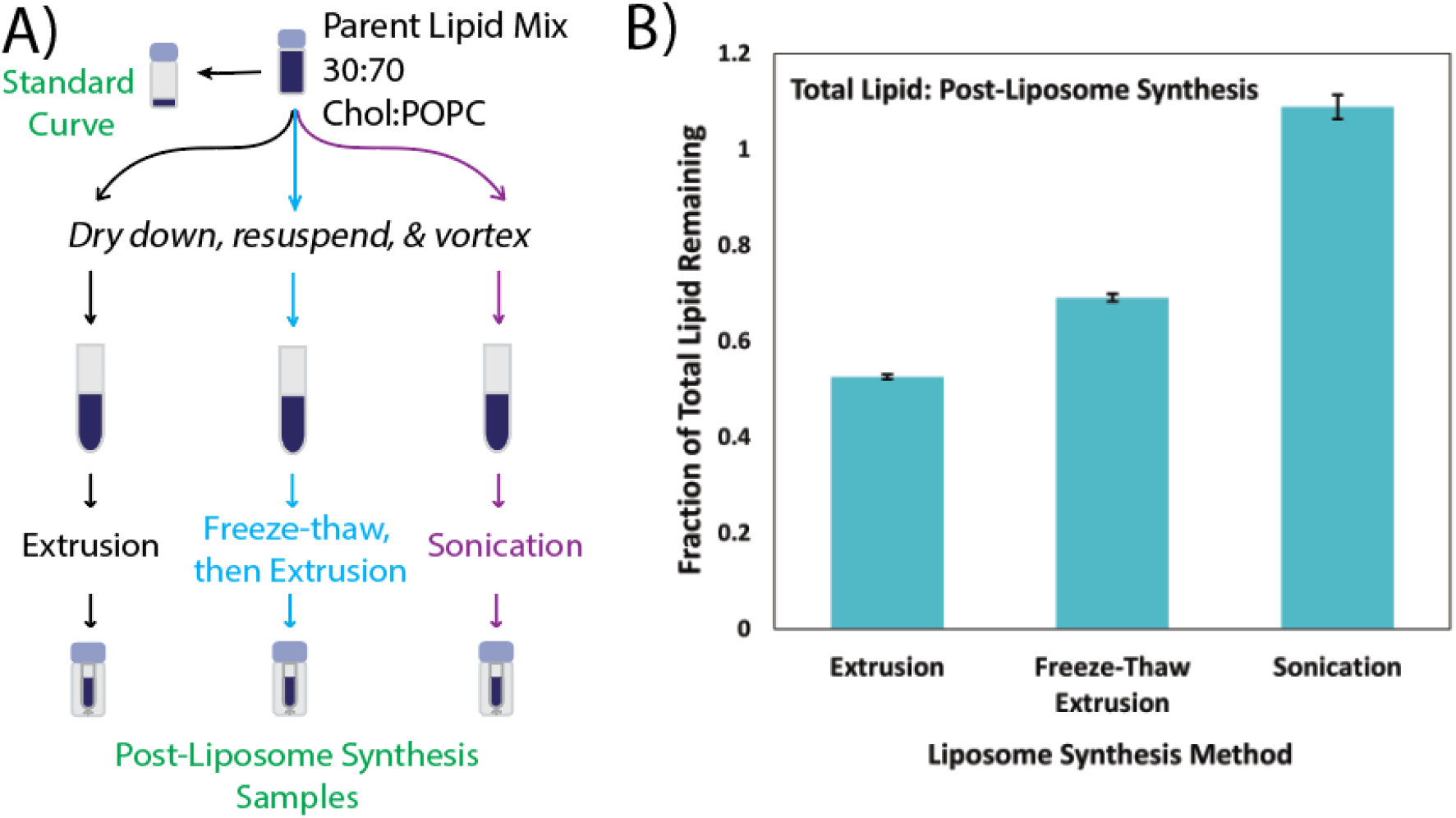
Fraction of total lipid remaining for liposomes prepared by extrusion, freeze-thaw extrusion, or sonication. **A)** shows a schematic of liposome preparation and sample collection for HPLC-ELSD analysis. All samples were prepared from 30:70 cholesterol: POPC lipid mixtures. Each post-liposome synthesis sample was taken immediately following the respective liposome synthesis method (see Materials and Methods Section 2.8) and analyzed by HPLC-ELSD. The Vortex Genie (30 seconds at max speed) was used to resuspend all samples. **B)** shows the fraction of total lipid remaining for each synthesis method, calculated relative to the theoretical maximum moles of lipid that could be recovered if the entirety of the starting lipid material ended up in that sample (see Equation 1). Data shown is mean ± propagated error of 3 measurements (see Equation 2 for error calculations).

We also used dynamic light scattering (DLS) to compare the distribution of liposome sizes produced by each of the 3 methods (see **Figure S2**). We observed that the Standard Extrusion and Freeze-Thaw Extrusion procedures yielded narrow distributions of liposome diameter (mean diameters = 136 nm and 126 nm respectively for liposomes extruded through 100 nm pore size). On the other hand, the Bath Sonication method produced a heterogeneous distribution with two peaks centered around 116 nm and 802 nm.

### 3.7. Influence of cholesterol mole fraction in lipid mixture

Various informal conversations we have had with researchers in the field, as well as some prior published research (26), suggest that preparing model membranes with high concentrations of cholesterol has the potential to lead to erroneous final lipid compositions, perhaps in part due to the maximum solubility limits of cholesterol which can vary artificially depending on preparation method (26). We examined this question directly in our Standard Extrusion procedure by preparing liposomes with cholesterol:POPC molar ratios of 30:70, 40:60, or 50:50 and quantifying lipid loss and compositional variance at each step (**Figure 6**).

**Figure 6.**
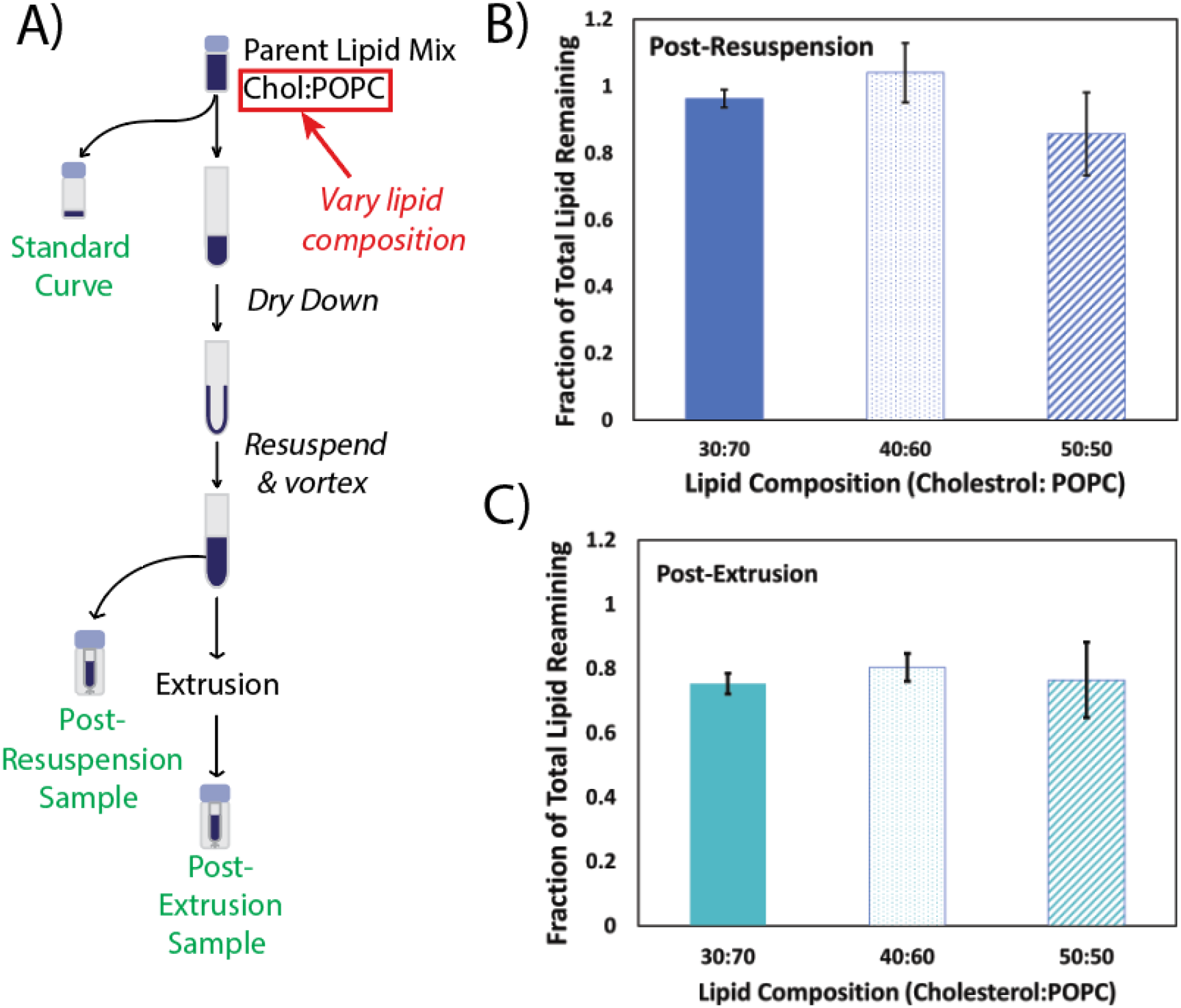
Fraction of total lipid remaining for post-resuspension and post-extrusion samples of liposomes prepared at different target lipid compositions. **A)** shows a schematic of liposome preparation and sample collection for HPLC-ELSD analysis. Liposomes were prepared by the Standard Extrusion Protocol at 3 different molar ratios of cholesterol:POPC = 30:70, 40:60, 50:50 (solid, dotted, striped data, respectively). Post-resuspension samples were taken immediately following the resuspension of the lipid film, but prior to extrusion. The BV1000 or Fisherbrand vortexer (30 seconds at max speed) was used to resuspend all samples. Post-extrusion samples were taken immediately following the extrusion step (see Methods & Materials, Section 2.8). All samples were analyzed by HPLC-ELSD. **B and C)** show the fraction of total lipid remaining for the post-resuspension and post-extrusion samples respectively. Each was calculated relative to the theoretical maximum moles of lipid that could be recovered if the entirety of the starting lipid material ended up in that sample (see Equation 1). Data shown is mean ± stdev of 2 or 3 replicate samples.

In the post-resuspension samples (**Figure 6b**), we observed that the 30:70 and 40:60 compositions exhibited essentially complete recovery of total lipid, whereas the 50:50 composition was somewhat lower; average fraction recovery was ∼0.85, with higher variability in the triplicate sample set.

In the post-extrusion samples (**Figure 6c)**, we observed that all compositions had lost additional lipid during the extrusion step, as expected (see **Section 3.5** above); final fraction recoveries were ∼0.7-0.8. However, the lipid composition was not altered substantially for any of the tested samples (**Figure 7**).

**Figure 7.**
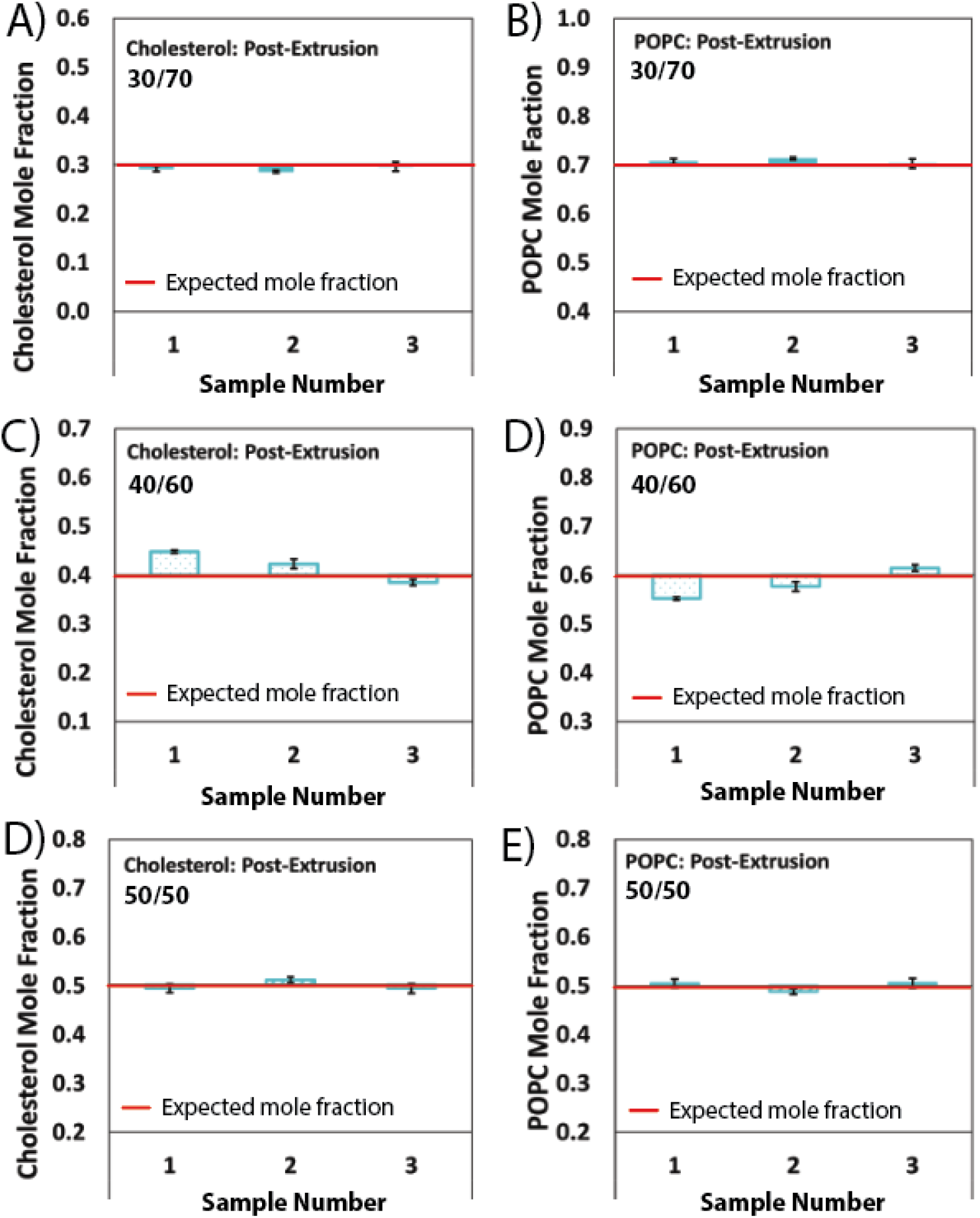
Measured mole fractions of cholesterol and POPC for post-extrusion samples of liposomes prepared at different lipid compositions. Liposomes were prepared by the Standard Extrusion Protocol at 3 different molar ratios of cholesterol:POPC = 30:70 **(A and B)**, 40:60 **(C and D)**, 50:50 **(E and F)**. The BV1000 vortexer (30 seconds at max speed) was used to resuspend all samples prior to extrusion. Post-extrusion samples were taken immediately following the extrusion step (see Methods & Materials Section 2.8, and schematic in Figure 6A) and analyzed by HPLC-ELSD. The expected mole fraction (red line) represents the mole fraction in the Parent Lipid Mixture prior to any manipulation. For each sample, data shown is the mean ± stdev of 2 or 3 replicate measurements.

Together, these data sets indicate that, in our hands, the Standard Extrusion procedure continues to produce reliable final lipid compositions up to 50:50 cholesterol:POPC, although with differing fractions of total lipid recovery at both the re-suspension and extrusion steps.

### 3.8. Best practices for reliable liposome preparation

Ideally, researchers would validate their own liposome preparation procedure using analytical methods such as the HPLC-ELSD approach we employ here. However, in the absence of such direct validation, we can offer suggestions for best practices based on our data. As noted above, we observed that the step with the largest impact on lipid loss was the re-suspension of the lipid film (**Figure 1**). However, if vigorous vortexing is utilized (minimum of 30 seconds on a newer vortex machine), maximum lipid recovery during re-suspension can reliably be achieved (**Figure 2**). Additionally, if extrusion is to be performed, freeze-thaw cycles prior to extrusion can be used as a secondary method to maximize lipid yield in the event the resuspension step doesn’t achieve complete lipid recovery (**Figure 5**). Sonication can also be employed to achieve the same ends (**Figure 5**). In our measurements, the extrusion step itself consistently results in ∼5-10% lipid loss, and so this should be accounted for by researchers who employ this method (**Section 3.5**) and are sensitive to that level of variation. Fortunately, in all the conditions we tested, only modest composition changes were observed, even when the overall lipid yield was low. It is possible however that such results may differ under other preparation conditions, and so researchers are recommended to validate their own approach as needed.

## 4. Conclusion

Using HPLC-ELSD, we studied lipid loss and compositional changes that can occur during preparation of simple 2-component liposomes, prepared by common methodologies. We found that the preparation methodology could have a substantial influence on lipid yield, with losses up to 50% or more depending on the method employed. The step with the largest impact was the re-suspension of the hydrated lipid film prior to liposome formation. Fortunately, if vigorous and lengthy vortexing is employed, reliably high lipid yields could be achieved. We note however that the common practice of vortexing until there is no visible residue on the bottom of the tube did not prove to be a reliable method. In many of the data sets discussed in this report, no obvious lipid residue was visible by eye even though substantial lipid losses were still detected by our HPLC-ELSD method. Both freeze-thaw and bath sonication methods can also improve lipid yields, more so than standard extrusion alone.

Under the conditions we tested, average compositional changes were modest at best, even in situations where the lipid loss was high. This should provide researchers some confidence in trusting the nominal lipid compositions in their own liposome preparations, but we note that it is possible that larger compositional changes may occur under conditions we did not test in this report. This underscores the importance of researchers assessing the lipid yield and compositional changes in their own liposome preparations, using analytical methods such as we have described herein.

## Author Contributions

EK, OG, RB, AL, and AB designed experiments, collected and analyzed data. EK and RB helped write the manuscript. RJR designed experiments, analyzed data, acquired project funding, and wrote the manuscript.

## Acknowledgments

The authors thank Dr. Nathan Cook and Dr. Silas Brown (Williams College) for instrument support. RJR acknowledges financial support from Williams College and NIH grant R15AI171754.

## Notes

### Competing Interest Statement

The authors have declared no competing interest.

