## Supplemental Information for "Lipid loss and compositional change during preparation of liposomes by common biophysical methods"

**Author Affiliations**
Department of Chemistry, Williams College, Williamstown, MA, 01267, USA

**Corresponding Author Contact Information**


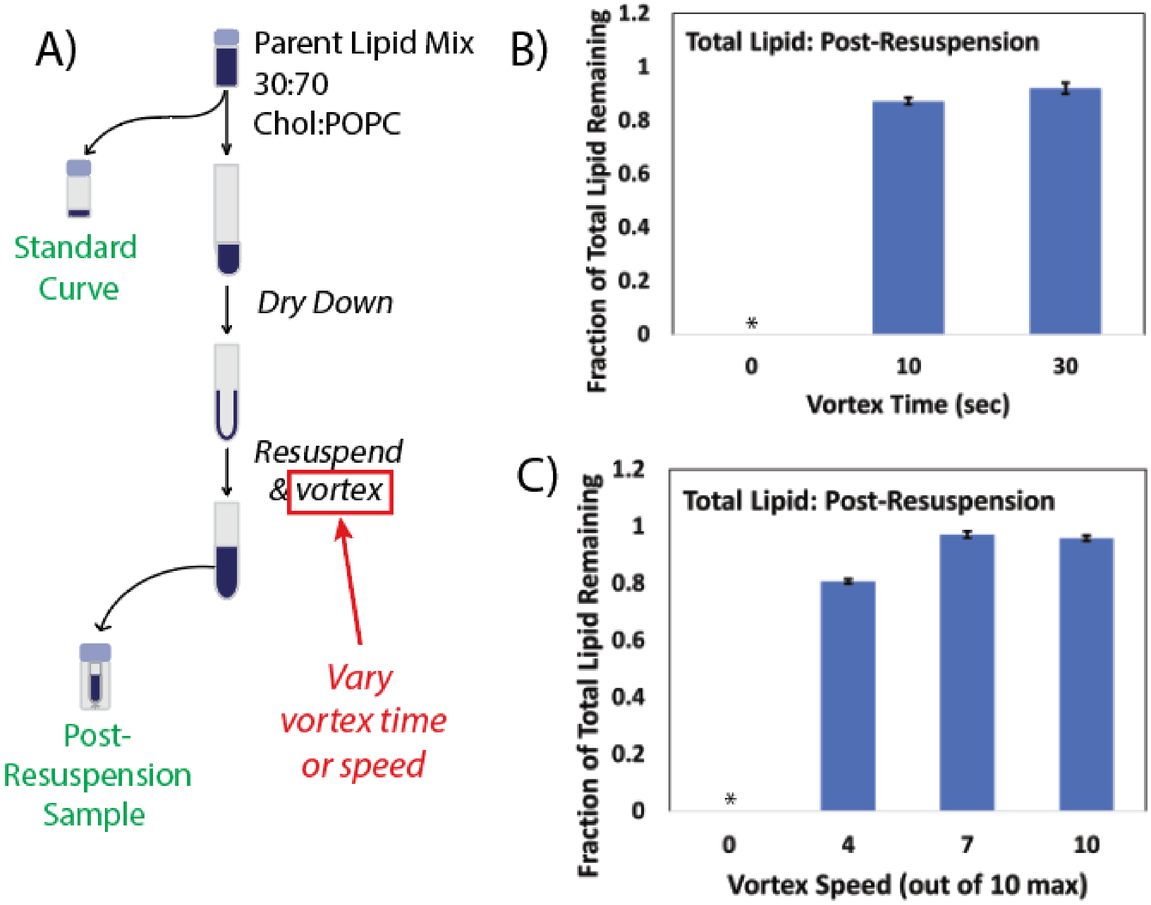


***Figure S1. Fraction of total lipid remaining for post-resuspension samples prepared by different vortexing times or speeds.*** ***A)*** *shows a schematic of sample preparation and collection for HPLC-ELSD analysis. Post-resuspension samples were prepared from 30:70 cholesterol: POPC lipid mixtures following the procedures listed in the Materials and Methods, but different vortex times or speeds were used to resuspend the lipid films. The Fisherbrand vortexer was used to resuspend all samples. Following sample collection, all samples were analyzed by HPLC-ELSD.* ***B)*** *shows the fraction of total lipid remaining for different vortex times.* ***C)*** *of total lipid remaining for different vortex speeds (max speed = 10, corresponding to 3200 rpm). Fraction of total lipid remaining was calculated relative to the theoretical maximum moles of lipid that could be recovered if the entirety of the starting lipid material ended up in that sample (see Equation 1 in the Main Text). Data shown is mean ± propagated error of 3 measurements (see Equation 2 in the Main Text for error calculations). * indicates that the ELSD signal fell below the lowest measurement on the standard curve, and was thus too low to be quantified.*

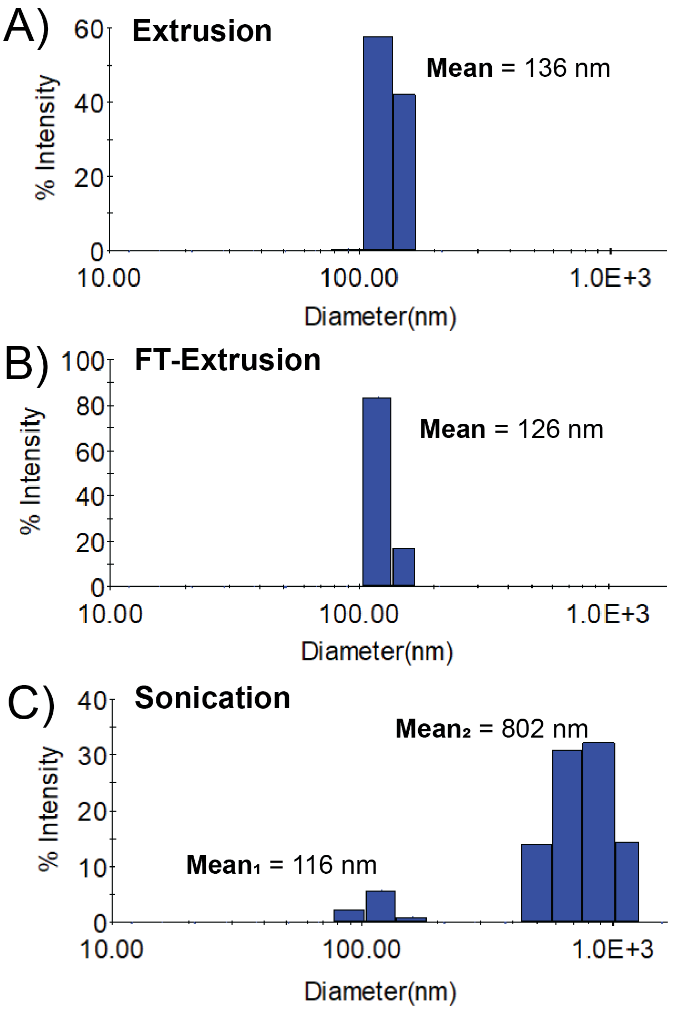


***Figure S2. Dynamic light scattering characterization of liposomes prepared by A) extrusion, B) freeze-thaw extrusion, and C) sonication.*** *Liposomes were composed of a nominal 70/30 mol% POPC/cholesterol ratio. Details on each preparation method are provided in the Materials and Methods. Whereas a single size peak was observed for liposomes prepared by extrusion and freeze-thaw extrusion (extrusion pore size = 100 nm), the sonication method produced multiple peaks. The mean diameter of each peak is noted.*
